## Supplementary material for "Non-*O* ABO blood group genotypes differ in their associations with *Plasmodium falciparum* rosetting and severe malaria"

**Table S1: *P. falciparum* parasite density by *ABO* genotype in severe malaria cases.**

| ***ABO genotype*** | ***Number of patients***^†^ | ***Parasite density/***μL **(95% CI)** | ***p* value** |
| --- | --- | --- | --- |
| *OO* | 612 | 62550 (50834 – 74265) | Reference |
| *AO* | 301 | 57406 (42096 – 72717) | 0.605 |
| *AA* | 37 | 58246 (14006 – 102486) | 0.858 |
| *BO* | 335 | 67952 (50723 – 85181) | 0.605 |
| *BB* | 33 | 72630 (14288 – 130972) | 0.723 |
| *AB* | 61 | 52147 (21218 – 83076) | 0.567 |
| *Non-O* | 767 | 61956 (51601 – 72310) | 0.994 |

Differences in parasite density by *ABO* genotype were tested by linear regression with adjustment for confounding by HbAS, ethnicity and gender. Comparisons were made between *ABO* genotype *OO* and individual non-*O* genotypes (*AO, AA, BO, BB & AB*) or all non-*O* genotypes combined.

^†^data were available for 1379 of the 1398 cases successfully genotyped for *ABO*. 17 cases had missing parasite density data and 2 had missing HbAS data.

**Table S2: Comparing Odds ratio differences for specific severe malaria syndromes between single dose and double dose non-*O* genotypes using the Wald test**

| **Case Phenotype** | ***ABO* genotype** | **No. of** | **Odds Ratio comparisons** | **Wald test P value** |
| --- | --- | --- | --- | --- |
|  |  | **Cases/controls** |  |  |
| *All CM* | *AO vs AA* | 166/810; 22/83 | 1.29/1.60 | 0.422 |
|  | *BO vs BB* | 160/683; 16/55 | 1.45/1.90 | 0.396 |
|  | *AO vs AB* | 166/810; 26/115 | 1.29/1.47 | 0.595 |
|  | *BO vs AB* | 160/683; 26/115 | 1.45/1.47 | 0.951 |
| *All SMA* | *AO vs AA* | 97/810; 9/83 | 1.35/1.18 | 0.736 |
|  | *BO vs BB* | 99/683; 10/55 | 1.62/1.71 | 0.896 |
|  | *AO vs AB* | 97/810; 19/115 | 1.35/2.05 | 0.137 |
|  | *BO vs AB* | 99/683; 19/115 | 1.62/2.05 | 0.394 |
| *All RD* | *AO vs AA* | 101/810; 13/83 | 1.41/1.68 | 0.598 |
|  | *BO vs BB* | 99/683; 20/55 | 1.63/2.01 | 0.586 |
|  | *AO vs AB* | 160/810; 12/115 | 1.41/1.25 | 0.715 |
|  | *BO vs AB* | 99/683; 12/115 | 1.63/1.25 | 0.414 |
| *Mortality* | *AO vs AA* | 33/810; 2/83 | 1.74/1.00 | 0.456 |
|  | *BO vs BB* | 23/683; 2/55 | 1.35/1.66 | 0.785 |
|  | *AO vs AB* | 33/810; 4/115 | 1.74/1.50 | 0.785 |
|  | *BO vs AB* | 23/683; 4/115 | 1.35/1.50 | 0.849 |

**Table S3: Cytoadhesion of *P. falciparum* line ItG by *ABO* genotype.**

| **ItG static adhesion to CD36** | | | | |
| --- | --- | --- | --- | --- |
| **N** | **Genotype** | **Mean pRBCs bound/mm^2^** | **95% CI** | **P value** |
| 51 | *OO* | 1385.64 | 1179.40 – 1591.89 | - |
| 32 | *AO* | 1345.61 | 1087.33 – 1603.89 | 0.814 |
| 6 | *AA* | 817.42 | 312.59 – 1322.25 | 0.073 |
| 17 | *BO* | 1146.66 | 813.49 – 1479.84 | 0.246 |
| 3 | *BB* | 1030.24 | 308.69 – 1751.79 | 0.386 |
| 3 | *AB* | 779.62 | 157.27 – 1401.96 | 0.123 |
| 61 | *Non-O* | 1191.53 | 1014.74 – 1368.32 | 0.163 |
| **ItG static adhesion to ICAM-1** | | | | |
| **N** | **Genotype** | **Mean pRBCs bound/mm^2^** | **95% CI** | **P value** |
| 51 | *OO* | 1640.72 | 1389.18– 1892.27 | - |
| 32 | *AO* | 1602.70 | 1286.15 – 1919.25 | 0. 854 |
| 6 | *AA* | 1567.11 | 772.38 – 2361.83 | 0.864 |
| 17 | *BO* | 1356.65 | 956.95 – 1756.34 | 0.254 |
| 3 | *BB* | 1296.59 | 398.11 – 2195.08 | 0.493 |
| 3 | *AB* | 1030.40 | 237.05 – 1823.75 | 0.208 |
| 61 | *Non-O* | 1479.30 | 1261.03 – 1697.57 | 0.340 |

Differences in ItG *P. falciparum* line binding to CD36 and ICAM-1 by *ABO* genotype were tested using multivariate linear regression analysis with adjustment for confounding by HbAS and α^+^thalassemia genotypes (including an interaction between HbAS and α^+^thalassemia). 112 RBC donor samples were tested once in duplicate over seven experimental days (day 1 n = 13, day 2 n = 8, day 3 n = 12, day 4 n = 1, day 5 n = 55, day 6 n = 15 and day 7 n = 8), therefore, experimental day was included as a co-variate to account for day-to-day variation.

**Table S4: ITvar9 PfEMP1 expression by *ABO* genotype.**

| **IT/R29 (ITvar9) PfEMP1 expression** | | | | |
| --- | --- | --- | --- | --- |
| **N** | **Genotype** | **Mean MFI** | **95% CI** | **P value** |
| 23 | *OO* | 3412.88 | 3180.61 – 3645.17 | - |
| 18 | *AO* | 3548.06 | 3283.57 – 3812.56 | 0.452 |
| 2 | *AA* | 3714.24 | 2869.41– 4559.07 | 0.487 |
| 9 | *BO* | 3325.30 | 2952.46– 3698.14 | 0.696 |
| 1 | *BB* | 3753.62 | 2617.84 – 4889.41 | 0.562 |
| 7 | *AB* | 3750.16 | 3317.22 – 4183.10 | 0.173 |
| 37 | *Non-O* | 3550.45 | 3372.44 – 3728.45 | 0.333 |
| **Percentage ITvar9 PfEMP1 expression** | | | | |
| **N** | **Genotype** | **% pRBCs ITvar9 positive** | **95% CI** | **P value** |
| 23 | *OO* | 56.71 | 53.74 – 59.69 | - |
| 18 | *AO* | 54.92 | 51.53 – 58.31 | 0.437 |
| 2 | *AA* | 57.94 | 47.13 – 68.76 | 0.824 |
| 9 | *BO* | 53.40 | 48.62 – 58.17 | 0.252 |
| 1 | *BB* | 56.40 | 41.86 – 70.95 | 0.967 |
| 7 | *AB* | 55.73 | 50.19– 61.28 | 0.756 |
| 37 | *Non-O* | 54.95 | 52.71 – 57.18 | 0.361 |

Differences in ITvar9 PfEMP1 expression by *ABO* genotype in the *P. falciparum* IT/R29 rosetting parasite line were tested by multivariate regression analysis with adjustment for confounding by HbAS and α^+^thalassemia (including interaction between HbAS and α^+^thalassemia). 60 RBC donor samples were tested once in duplicate over two successive experimental days (day 1 n = 30 and day 2 n = 30), therefore, experimental day was included as a co-variate to account for day-to-day variation.

**Table S5: General characteristics of the Kilifi longitudinal cohort study by *ABO* genotype.**

| **Characteristics** |  | ***OO (%)*** | ***AO (%)*** | ***AA (%)*** | ***BO (%)*** | ***BB (%)*** | ***AB (%)*** | ***p value*** |
| --- | --- | --- | --- | --- | --- | --- | --- | --- |
| **Sample size** | n = 242 | 135 (55.8) | 52 (21.5) | 2 (0.8) | 44 (18.2) | 4 (1.6) | 5 (2.1) |  |
| **Gender** | Males | 58 (56.3) | 21 (20.4) | 0 (0.0) | 20 (19.4) | 2 (1.94) | 2 (1.94) |  |
|  | Females | 77 (55.4) | 31 (22.3) | 2 (1.4) | 24 (17.3) | 2 (1.4) | 3 (2.2) | 0.933 |
| **Ethnic group** | Giriama | 118 (56.5) | 44 (21.1) | 1 (0.5) | 39 (18.6) | 4 (1.9) | 3 (1.4) |  |
|  | Chonyi | 8 (38.1) | 6 (28.5) | 1 (4.8) | 5 (23.8) | 0 (0.0) | 1 (4.8) |  |
|  | Others | 9 (75.0) | 2 (16.7) | 0 (0.0) | 0 (0.0) | 0 (0.0) | 1 (8.3) | 0.120 |
| **Sickle** | AA | 124 (58.5) | 43 (20.3) | 2 (0.9) | 38 (17.9) | 2 (0.9) | 3 (1.4) |  |
|  | AS | 11 (36.6) | 9 (30.0) | 0 (0.0) | 6 (20.0) | 2 (6.67) | 2 (6.67) | 0.029 |
| **α+thalassaemia** | αα/αα | 41 (55.4) | 20 (27.0) | 1 (1.4) | 11 (14.8) | 1 (1.4) | 0 (0.0) |  |
|  | -α/αα | 75 (61.5) | 16 (13.1) | 1 (0.8) | 25 (20.5) | 2 (1.6) | 3 (2.5) |  |
|  | -α/-α | 19 (41.3) | 16 (34.7) | 0 (0.0) | 8 (17.4) | 1 (2.2) | 2 (4.4) | 0.043 |
| **Age in months**  **Median (IQR)** |  | 32.2 (16-49) | 30 (12-50) | 39 (30-49) | 39 (20-53) | 40 (18-54) | 39 (18-49) | <0.001 |

The Fisher’s exact test was used to test for differences in the distribution of *ABO* genotypes across categorical variables of gender, ethnic group, HbS and α^+^thalassemia genotypes while the Kruskal-Wallis test was used to test for differences in age (as a continuous variable) by *ABO* genotype. IQR, interquartile range.

**Table S6: Incidence rate ratios (IRR) for uncomplicated malaria in Kenya by *ABO* genotype**

|  |  |  | **Crude** | | | |  | **Adjusted**^†^ | | | |
| --- | --- | --- | --- | --- | --- | --- | --- | --- | --- | --- | --- |
| **N** | **Episodes** | ***ABO* genotype** | **IRR** | **LCI** | **UCI** | **P** |  | **IRR** | **LCI** | **UCI** | **P** |
| 135 | 528 | *OO* | 1 |  |  |  |  | 1 |  |  |  |
| 52 | 237 | *AO* | 1.13 | 0.80 | 1.60 | 0.495 |  | 1.26 | 0.90 | 1.78 | 0.181 |
| 2 | 10 | *AA* | 1.19 | 0.27 | 5.31 | 0.817 |  | 1.03 | 0.26 | 4.05 | 0.963 |
| 5 | 19 | *AB* | 0.81 | 0.30 | 2.16 | 0.672 |  | 1.01 | 0.40 | 2.56 | 0.986 |
| 44 | 158 | *BO* | 0.81 | 0.56 | 1.18 | 0.279 |  | 0.95 | 0.66 | 1.37 | 0.787 |
| 4 | 6 | *BB* | 0.37 | 0.10 | 1.38 | 0.139 |  | 0.58 | 0.16 | 2.11 | 0.411 |
| 107 | 430 | *Non-O** | 0.96 | 0.72 | 1.27 | 0.753 |  | 1.09 | 0.83 | 1.45 | 0.532 |

Incidence rate ratios and 95% confidence intervals were generated using a random effects Poisson regression analysis, either without or ^†^with adjustment for age, season, ethnic group, HbAS and α^+^thalassaemia. The analysis also took into account within person clustering of events. Data represent 250, 88.1, 4.5, 10.7, 82.6, & 6.1 child years of follow up for *OO,* *AO, AA, AB, BO* and *BB* genotypes respectively. * Analysis comparing non-*O* to *OO* was done using a recessive model of inheritance.

LCI: lower 95% confidence interval; UCI: upper 95% confidence interval.

**Table S7: Odds ratios (OR) for asymptomatic malaria in Kenya by *ABO* genotype.**

|  |  |  | **Crude** | | | |  | **Adjusted**^†^ | | | |
| --- | --- | --- | --- | --- | --- | --- | --- | --- | --- | --- | --- |
| **N** | **Prevalence^§^** | ***ABO* genotype** | **OR** | **LCI** | **UCI** | **P** |  | **OR** | **LCI** | **UCI** | **P** |
| 124 | 44/360 (12%) | *OO* | 1 |  |  |  |  | 1 |  |  |  |
| 48 | 24/130 (18.5%) | *AO* | 1.63 | 0.90 | 2.97 | 0.110 |  | 1.84 | 1.00 | 3.38 | 0.051 |
| 2 | 0/7 (0.0%) | *AA* | - | - | - | - | - | - | - | - | - |
| 5 | 3/16 (18.8%) | *AB* | 1.66 | 0.20 | 13.52 | 0.635 |  | 1.77 | 0.20 | 15.86 | 0.749 |
| 43 | 18/127 (14.2%) | *BO* | 1.19 | 0.62 | 2.27 | 0.598 |  | 1.12 | 0.56 | 2.26 | 0.743 |
| 2 | 2/7 (28.6%) | *BB* | 2.88 | 0.52 | 15.94 | 0.225 |  | 3.34 | 0.66 | 16.91 | 0.146 |
| 100 | 47/287 (16.4%) | *Non-O** | 1.41 | 0.84 | 2.36 | 0.190 |  | 1.45 | 0.85 | 2.46 | 0.170 |

^§^Prevalence: Number of *P. falciparum* positive slides/total slides (%). These data were derived from four cross-sectional surveys as part of the Kilifi longitudinal cohort study carried out in March, July and October 2000 and June 2001. Odds ratios and 95% confidence intervals were generated using a logistic regression analysis, without or ^†^with adjustment for age, season, ethnic group and HbAS genotype. The analysis also took into account within person clustering of events. *Analysis comparing non-O to blood group O was done using a recessive model of inheritance.

Abbreviations: OR, odds ratio (95% confidence interval). LCI: lower 95% confidence interval; UCI: upper 95% confidence interval.

**Supplementary figure legends**

**Figure S1:** *P. falciparum* IT/R29 ABO blood group rosetting preference.

ABO blood group preference assay of the IT/R29 parasite strain was assessed using fluorescently-labelled RBC from 13 donors (4 group O, 4 group A and 5 group B). “Home O” indicates the blood group O donor that was used to culture the parasites. The y-axis shows the difference between the percentage of labelled cells found in rosettes, compared to the percentage found in the mix. Experiments were carried out in triplicate, and the mean and SEM are shown for each donor. For statistical analysis, the triplicate values for each donor were averaged and treated as a single data point, such that n=4 for groups O and A, and n=5 for group B. The blood groups were compared using a Kruskal Wallis test with Dunn’s multiple comparisons (** P<0.01).

**Figure S2.** Relative cytoadherence of *P. falciparum* ItG by *ABO* genotype.

**(a)** Relative binding to CD36 recombinant protein. **(b)** Relative binding to ICAM-1 recombinant protein. Purified ITg infected RBCs (iRBCs) were allowed to invade into RBCs from 112 donors (*OO* n=51, *AO* n=32, *AA* n=6, *BO* n=17, *BB* n=3, *AB* n=3) and static adhesion to immobilized recombinant proteins was tested the next day. Samples were tested over seven experimental days (day 1 n = 13, day 2 n = 8, day 3 n = 12, day 4 n = 1, day 5 n = 55, day 6 n = 15 and day 7 n = 8), with each donor being tested once. For each RBC sample, adhesion was tested in two dishes with triplicate protein spots in each dish and the data presented as the mean iRBC bound/mm^2^ for each donor. Because baseline binding using a single donor varies from day to day, the binding data for each sample were normalized to that of the mean binding for the control iRBCs (*OO*) run on the same day (number of reference *OO* genotype samples run each day*;* day 1 n = 7, day 2 n = 4, day 3 n = 8, day 4 n =1, day 5 n = 23, day 6 n = 7 and day 7 n = 2). Horizontal bars represent median relative adhesion for each genotype. Number of samples per genotype are shown in parenthesis. A Kruskal-Wallis test gave P=0.310 for CD36 adhesion and P=0.814 for ICAM-1 adhesion.

**Figure S3.** Relative PfEMP1 expression in *P. falciparum* IT/R29 by *ABO* genotype.

**(a)** Relative ITvar9 PfEMP1 expression. **(b)** Relative ITvar9 positive iRBCs. Purified IT/R29 infected RBCs (iRBCs) were allowed to invade into RBCs from 60 donors (*OO* n=23, *AO* n=18, *AA* n=2, *BO* n=9, *BB* n=1, *AB* n=7) and PfEMP1 expression was assessed the next day by flow cytometry with antibodies specific for the ITvar9 PfEMP1 variant. Samples were tested over two consecutive experimental days (day 1=30 and day 2=30). Median fluorescent intensity and proportion of ITvar9 positive iRBC data for all samples were normalized to that of the mean MFI and proportion of ITvar9 positive iRBCs for the control pRBCs (*OO*) (14 and 9 reference *OO* genotype samples run on day 1 and 2 respectively) run on the same day. Horizontal bars represent median relative MFI and proportion of ITvar9 positive iRBCs for each genotype. Kruskal-Wallis test P=0.076 for MFI and P=0.285 for % positive iRBCs.

**Figure S1**

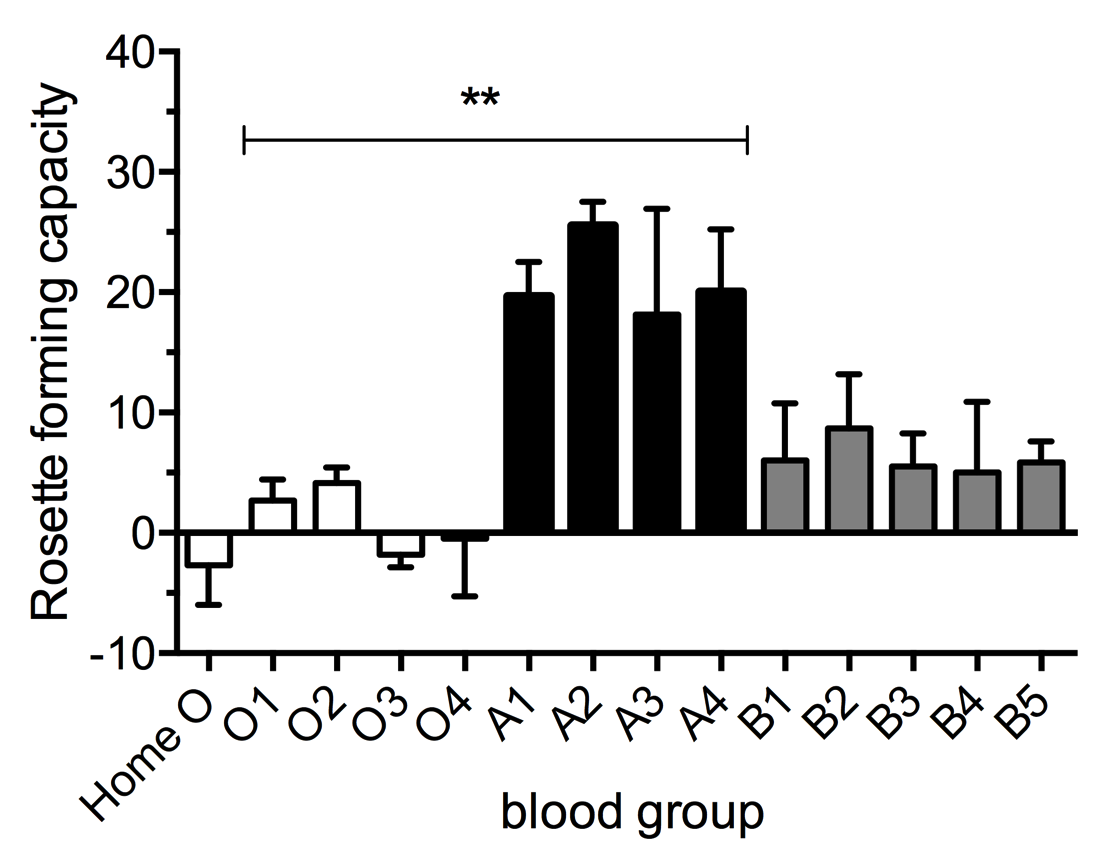

**Figure S2**

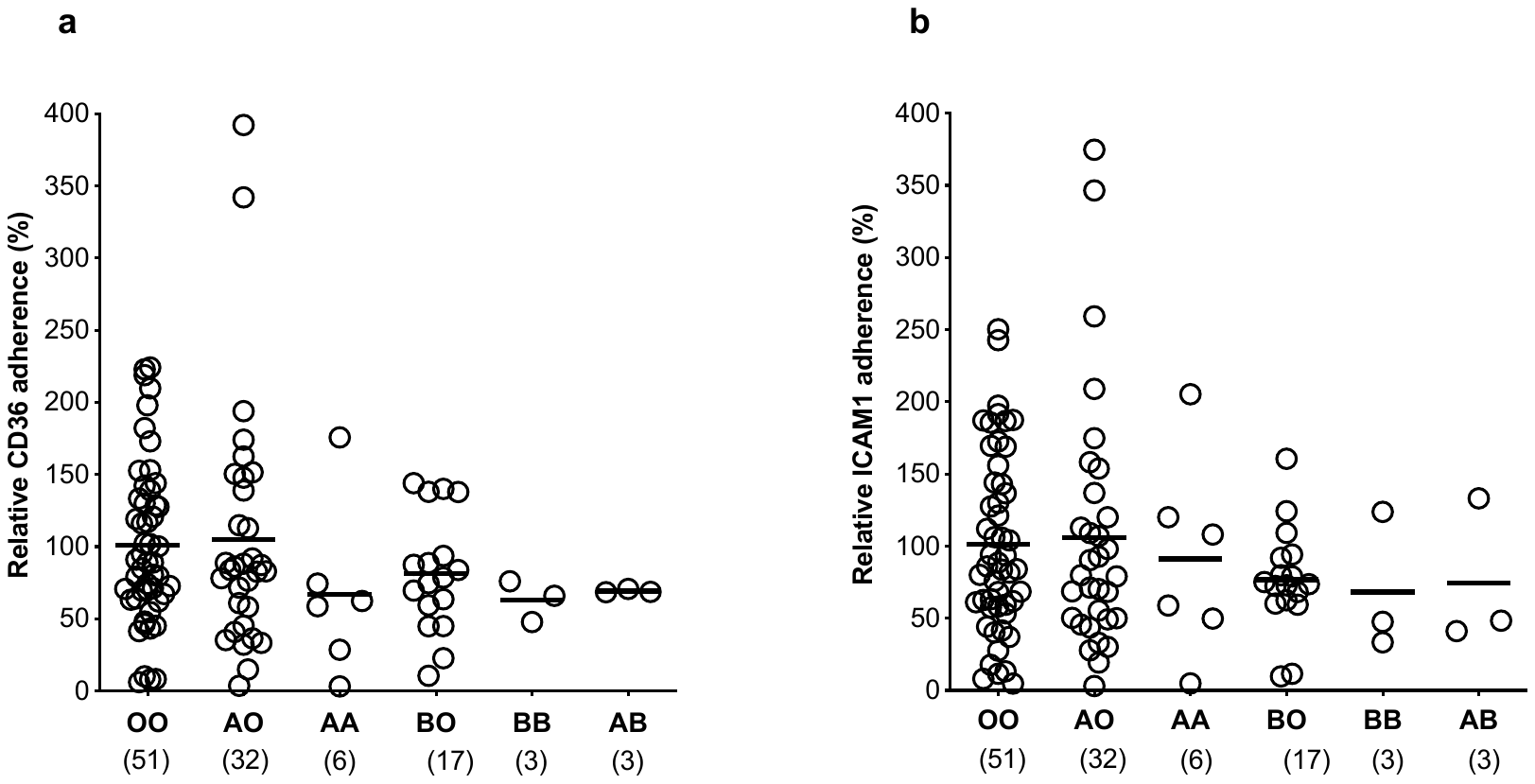

**Figure S3**

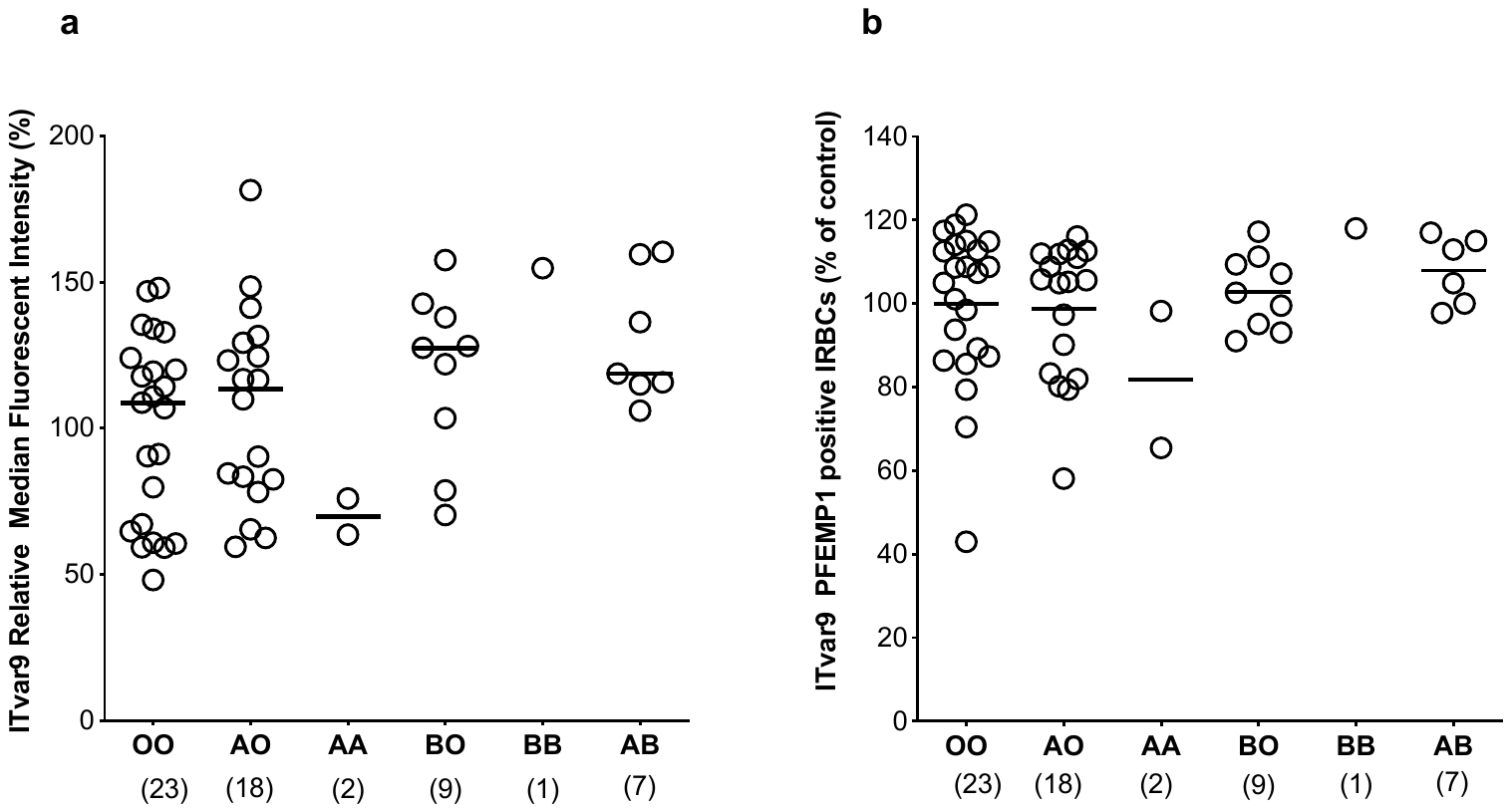
